# Kif6 regulates cilia motility and polarity in brain ependymal cells

**DOI:** 10.1101/2023.02.15.528715

**Authors:** Maki Takagishi, Yang Yue, Ryan S. Gray, Kristen J. Verhey, John B. Wallingford

## Abstract

Ependymal cells, lining brain ventricular walls, display tufts of cilia that beat in concert promoting laminar Cerebrospinal fluid (CSF) flow within brain ventricles. The ciliary axonemes of multiciliated ependymal cells display a 9+2 microtubule array common to motile cilia. Dyneins and kinesins are ATPase microtubule motor proteins that promote the rhythmic beating of cilia axonemes. Despite common consensus about the importance of axonemal dynein motor proteins, little is known about how Kinesin motors contribute to cilia motility. Here, we define the function of Kinesin family member 6 (Kif6) using a mutation that lacks a highly conserved C-terminal tail domain (*Kif6^p.G555fs^*) and which displays progressive hydrocephalus in mice. An analogous mutation was isolated in a proband displaying macrocephaly, hypotonia, and seizures implicating an evolutionarily conserved function for Kif6 in neurodevelopment. We find that loss of Kif6 function caused decreased ependymal cilia motility and subsequently decreased fluid flow on the surface of brain ventricular walls. Kif6 protein was localized at ependymal cilia and displayed processive motor movement (676 nm/s) on microtubules *in vitro*. Loss of the Kif6 C-terminal tail domain did not affect the initial ciliogenesis *in vivo*, but did result in defects in cilia orientation, the formation of robust apical actin networks, and stabilization of basal bodies at the apical surface. This suggests a novel role for the Kif6 motor in maintenance of ciliary homeostasis of ependymal cells.

**Summary statement:** We found that Kif6 is localized to the axonemes of ependymal cells. In vitro analysis shows that Kif6 moves on microtubules and that its loss mice decrease cilia motility and cilia-driven flow, resulting in hydrocephalus.

## Introduction

Multiciliated cells have dozens of motile cilia, which are microtubule-based projections. These motile cilia are planar polarized and coordinately beat to generate directional flow across an epithelium (Brooks and Wallingford, 2014). Ciliary-driven directional fluid flow maintains the cerebrospinal fluid (CSF) circulation in the brain, mucus clearance in the respiratory tract, and the transport of gametes in the reproductive system (Spassky and Meunier, 2017).

Dyneins and kinesins are ATPase microtubule motor proteins that drive diverse motion in cilia and flagella (Gennerich and Vale, 2009). For example, dynein complexes, such as outer (ODA) and inner (IDA) dynein arms are arrayed along microtubule doublets to generate motive force for cilia beating (Ishikawa, 2017). In addition, both kinesins and dyneins help facilitate intraflagellar transport of cargoes necessary to build and maintain cilia (Prevo et al., 2017).

Kinesin-9 family members Kif6 and Kif9 are both implicated in cilia function. For example, the *Trypanosome brucei* Kinesin-9 family members, KIF9A and KIF9B were localized in the axoneme and their knockdown reduced flagellum motility (Demonchy et al., 2009). The KIF9A homologues *Chlamydomonas reinhardtii* KLP1 and *human/mouse* Kif9 were recently shown to be active motors that act in the central pair to facilitate ciliary beating in *Chlamydomonas* and vertebrates (Lechtreck and Witman, 2007; Konjikusic et al., 2023). Less is known about the KIF9B homologue *human/mouse* Kif6. In humans, multiple studies showed an association between the *KIF6* 719Arg allele and an increased risk of cardiovascular disease (Bare et at., 2007; Shiffman et al., 2008a; Shiffman et al., 2008b; Li et al., 2018). However, a subsequent meta-analysis of multiple case-control studies failed to find evidence of this association (Assimes et al., 2010).

In contrast, deleterious mutations in *KIF6* are associated with neurodevelopmental defects in patients, including seizures, hypotonia, and macrocephalus (Gray et al., 2021; Konjikusic et al., 2018). In animal models, loss of Kif6 function resulted in hydrocephalus and decreased cilia numbers in the adult brain ventricles of both mice and zebrafish (Gray et al., 2021; Konjikusic et al., 2018). Despite these clinical physiological phenotypes, the molecular function of Kif6, remains unresolved.

Here, we provide evidence that Kif6 functions as a canonical kinesin motor undergoing processive microtubule-based movement and localizes in the motile cilia and cytoplasm of mouse brain ependymal cells. Loss of the Kif6 elicits a complex array of cilia-related phenotypes marked by defects in cilia beating, CSF fluid flow, and planar polarization of basal bodies. Over time, ependymal cilia in Kif6 mutant mice are lost, possibly due to defects in apical actin assembly and subsequent destabilization of basal bodies. Thus, our data suggest that Kif6 plays several roles in ependymal cell cilia homeostasis and neurodevelopment in mammals and may shed light on Kif6-related human pathologies.

## Results

### Kif6 is an active motor that localizes to cytoplasmic and ciliary microtubules in ependymal cells

We previously reported a macrocephaly patient with a mutation in *KIF6* that lacks part of the stalk region and C-terminal tail (Konjikusic et al., 2018), which is the putative cargo-binding domain (Piddini et al., 2001). We also generated mice with a similar mutation lacking its C-terminal stalk and tail domains, which displayed variably penetrant hydrocephalus and reduction in ependymal cell cilia (Konjikusic et al., 2018). Since the motor activity of Kif6 has never been determined, these findings led us to perform in vitro studies.

The alignment of the protein sequences of human (h), mouse (m), and *Danio rerio (d)* Kif6 proteins showed that the overall domain organization is highly conserved (Fig.1A). When overexpressed in COS-7 cells, full-length (FL) dKif6-EGFP did not localize to microtubules but rather showed a diffuse localization within the cytosol (Fig. S1). In *in vitro* single-molecule motility assays, dKif6 (FL)-EGFP did not bind to microtubules (Fig. S1), suggesting that full-length dKif6 is regulated by auto-inhibition, which is common for kinesin proteins (Verhey and Hammond, 2009). To examine the motility properties of dKif6, we generated a truncated version, dKif6(1-501)-EGFP, which includes the motor domain and the first two coiled coil segments for homodimerization (Fig. 1A). When overexpressed in COS-7 cells, dKif6 (1-501)-EGFP accumulated at the cell periphery, suggesting that dKif6 (1-501) is capable of movement to the microtubule plus ends, although a large fraction of protein showed diffuse localization within the cytosol (Fig. 1B). To visualize Kif6 movement directly, we carried out *in vitro* single-molecule motility assays and found that dKif6 (1-501)-EGFP moved processively along microtubules with a speed of 676.7 ± 257.6 nm/sec (Mean ± SD) (Fig.1C-E, Movie 1), suggesting that Kif6 could serve as a transporter of axonemal components in ependymal cell cilia.

**Figure 1.**
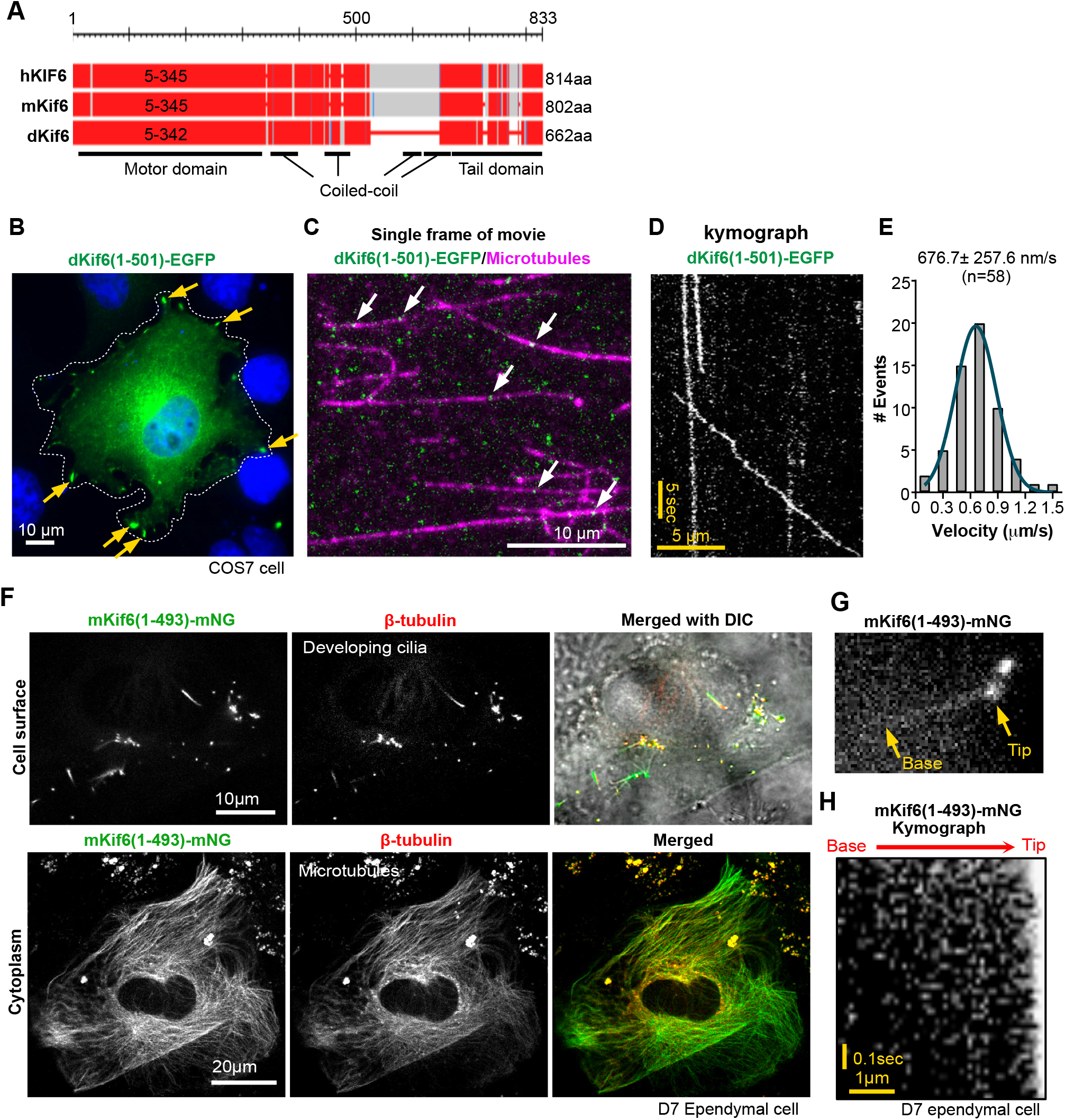
Kif6 is an active motor that localizes to cytoplasmic and ciliary microtubules. (A) Schematic representation of human, mouse, and danio Kif6 proteins. The predicted motor domain is 5-345aa in hKIF6, 5-345aa in mKif6, and 5-342aa indKif6. Red regions are highly conserved positions with human, mouse, and danio. (B) Representative image of GFP-tagged dKif6(1-501) expressed in COS-7 cells. White dashed line indicates the outline of a transfected cell. Yellow arrowheads indicate the accumulation of dKif6(1-501)-EGFP at microtubule plus ends in the cell periphery. Blue, DAPI stain. (C) Representative still image from single-molecule motility assay of dKif6(1-501)-EGFP molecules (green) on taxol-stabilized microtubules (magenta). White arrowheads indicate individual dKif6(1-501)-EGFP molecules on microtubules. Scale bar, 10 μm. (D) Kymograph of dKif6(1-501)-EGFP movement from the single-molecule motility assay shown in (C). Time is on the y-axis (scale bar, 5 sec) and distance is on the x-axis (scale bar, 5 μm). (E) The velocities of individual dKif6(1-501)-EGFP were determined from kymographs and plotted as a histogram for the population. The curve was fit with a normal distribution. Velocity is described as mean ± std. dev. n=58 motility events analyzed across three independent experiments. (F) mKif6(1-493)-mNeongreen (NG) expressed in cultured developing ependymal cell at D7. Upper panels show mKif6(1-493)-mNG (green), immunostained-β-tubulin (red) and merged image with DIC on cell surface to see cilia. Lower panels show mKif6(1-493)-mNG and β-tubulin (red) in the same cell at the cytoplasmic plane. (G) Still frame from live imaging of mKif6 (1-493)-mNG in a developing short cilium of a D7 ependymal cell. Arrows indicate ciliary base and tip. The upper dot is the tip of another cilium. See supplemental movie 2. (H) Kymograph of GFP-mKif6 (1-493) movement along cilia in the live imaging shown in (F).

A similarly truncated version of mouse Kif6, mKif6 (1-493)-mNeonGreen (mNG), was found to co-localize with β-tubulin in short, developing cilia as well as along cytoplasmic microtubules in cultured ependymal cells at 7 days after differentiation (D7) (Fig. 1F). mKif6 (1-493) accumulated at the tip of short cilia (Fig. 1G), consistent with microtubule-based processive movement, however, we were unable to observe any clear movement of mKif6 (1-493) within the cilia (Fig. 1H, Movie 2). Together these data suggest that the Kif6 motor domain derived from both zebrafish and mice is able to bind to and move processively along microtubules.

We next set out to define endogenous Kif6 protein localization using immunochemistry. Antibody specificity was confirmed using Western blot analysis of primary ependymal cell lysates derived from WT and *Kif6^pG555f/pG555f^* homozygous mutant mice (hereafter labeled *Kif6^pG555f^*). The *Kif6^pG555fs^* mutation is predicted to truncate Kif6 protein thus removing the antigen region of the Kif6 antibody. We confirmed a single band of Kif6 protein expression in testis and in primary cultured ependymal cells by Western blot (Fig 2A). Interestingly, we consistently observed multiple bands using the Kif6 antibody in lysates derived from Lateral Ventricular (LV) wall tissues. Because LV wall tissues contains many cells other than pure populations of primary cultured ependymal cells, robust Kif6 expression may be masked in these lysates. Next we enriched for ependyma from primary LV tissue and confirmed ependymal cell differentiation at D0, D5, and D14. At D0, the precursor cells had Arl13b-positive a short cilium (primary cilia). At D5, the cultured dish included precursor cells and immature ependymal cells that had short or elongating multiple cilia. At D14, mature ependymal cells had multiple long cilia (Fig. 2B). We first observed Kif6 at D5 of differentiation, which is concomitant with the onset of multiciliated cell differentiation in cultured ependymal cells.

**Figure 2.**
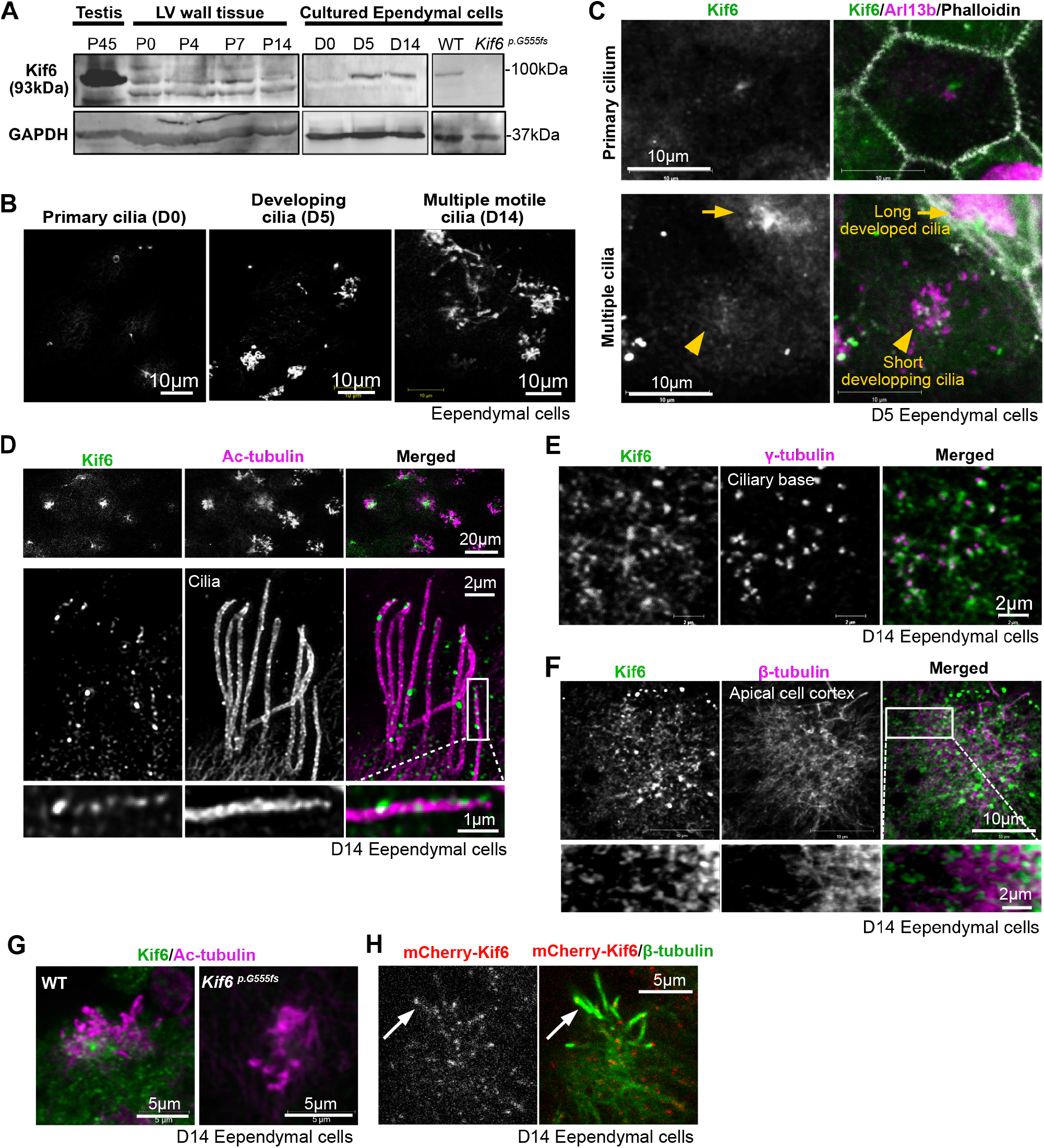
Kif6 localizes to cilia and cytoplasmic microtubules in ependymal cells. (A) Western blot analysis with anti-Kif6 or anti-GAPDH in P45 mouse testis tissue, P0, 4, 7, or 14 mouse LV wall tissues, or D0, 5, or 14 cultured ependymal cell lysates. Right panels show D14 ependymal cell lysate from WT or *Kif6^p.G555fs^* mice. (B) Representative image of cilia formation in cultured ependymal cells at D0, 5, or 14. ependymal cells were immunostained with anti-Arl13b. (C) Immunostainning with anti-Kif6 (green), anti-acetylated (Ac)-tubulin (magenta, as a cilia marker), and Phalloidin (gray, as a cell border marker) in differentiating ependymal cells at D5. Upper panels show primary ciliated precursor cell. Lower panels show multiple ciliated differentiating ependymal cells. (D) Super-resolution image of immunostaining with anti-Kif6 (green) and anti-Acetylate (Ac)-tubulin (magenta) in differentiated ependymal cell cilia at D14. Top panels show Kif6 expression in the differentiated ependymal cells at low magnification. Middle panels show representative ependymal cilia in one ependymal cell. Boxed area is enlarged in the bottom panels. (E) Immunostaining with anti-Kif6 (green) and anti-γ-tubulin (magenta, as a BF marker) at the cytoplasmic ciliary base. (F) Immunostaining with anti-Kif6 (green) and anti-β-tubulin (magenta, as a microtubules marker) at the apical cell cortex. (G) Immunosatining with anti-Kif6 (green) at cilia and ciliary base in D14 ependymal cell from WT or *Kif6^p.G555fs^* as negative control. (H) mCherry-Kif6 (white in left panel, red in right panel) expressing cells are immunostained with anti-β-tubulin (green). Arrow indicates mCherry-Kif6 localization at cilia.

We next examined Kif6 localization in ependymal cells by immunostaining *in vivo*. At D5 we observed Kif6 punctum close to primary cilium, with increased accumulation at the base of both short developing and long developed multicilia (Fig. 2C). In order to resolve these accumulations of Kif6 in multicilia we used super-resolution imaging in D14 ependymal cell cultures, showing puncta along the length of ciliary axonemes (Fig. 2D). We also observed Kif6 puncta at the apical cell cortex localized with the basal foot marker, γ-tubulin (Fig. 2E), and partially co-localized with microtubules (β-tubulin) at apical cell cortex around the ciliary base (Fig. 2F). The specificity of immunostaining with Kif6 antibody was confirmed by the loss of signal at multicilia in D14 ependymal cells derived from *Kif6^pG555fs^* mutant mice (Fig. 2G). Finally, an exogenously expressed N-terminal fluorescently tagged mCherry-Kif6 protein localized at multicilia and with apical β-tubulin (Fig. 2H). These results suggest that endogenous Kif6 localizes at the apical tubulin cortex, at the base of cilia, and within ciliary axonemes.

### Kif6 regulates cilia motility in ependymal cells

As noted, *Kif6^pG555fs^* mice display variably penetrant hydrocephalus (Fig. 3). We observed moderate hydrocephalus with enlarged LV in 69.2% (9/13 mice) while severe hydrocephalus, often associated with broken LV and an absence of ependymal cells, was observed in the remainder (Fig. 3A). Ependymal cilia were present in the middle part of LV wall in *Kif6^pG555fs^* mice with moderate hydrocephalus (Fig. 3B-C), so we analyzed these cells at P7-14 for observation of cilia phenotypes.

**Figure 3.**
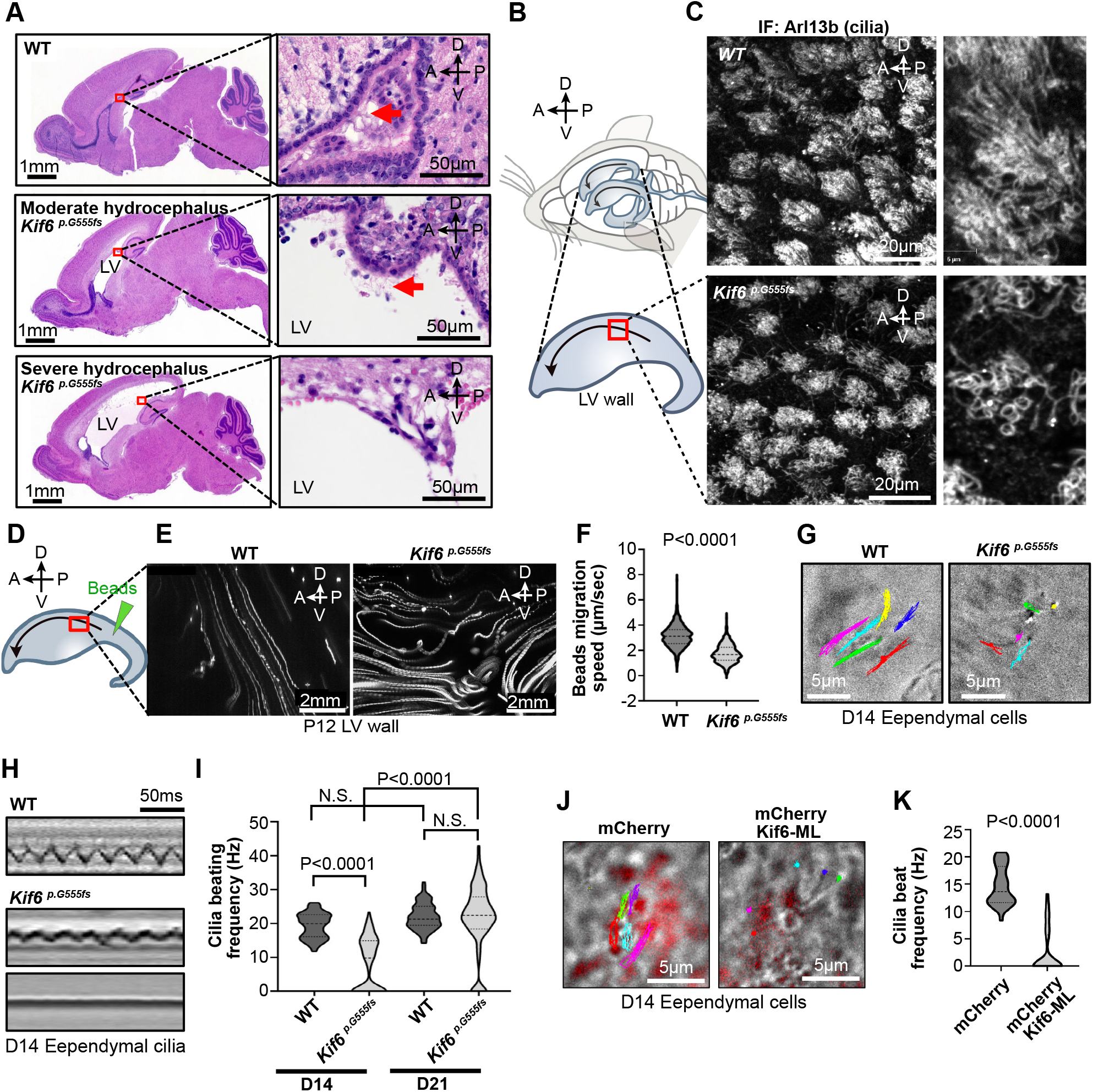
Kif6 regulates cilia motility in ependymal cells. (A) H&E-stained sagittal section in WT or *Kif6^p.G555fs^* mice brains at P14. Red boxes indicate the wall of Lateral ventricle (LV) expanded in the right panels. Arrows represent ependymal cilia. LV wall in *Kif6^p.G555fs^* mice with severe hydrocephalus is broken and lose ependymal cells. (B) Illustration of mouse brain ventricles and dissected LV wall. Arrows represent CSF flow direction. CSF is produced at the choroid plexus in the posterior region within LV, and outflows through the Foremen of Monro at the anterior-ventral region of the LV, towards the third ventricle. The illustration below represents the surface of a dissected right distal LV wall. A, anterior, P, posterior, D, dorsal, V, ventral sides of the brain. (C) Fluorescence microscopy images of the red boxed area in (C) are shown to see the surface cilia formation in P14 WT or *Kif6^p.G555fs^*. LV wall tissues were whole-mount immunostained with anti-Arl13b as a cilia marker. (D) Representation of beads migration assay on the surface of LV wall tissue at P12. Fluorescent microbeads are placed on the green arrowhead region and migrate toward the anterior-ventral side along the CSF flow (arrow). (E) Time projections of beads migration on WT or *Kif6^p.G555fs^* LV wall at the red boxed area shown in (E). 100 flames are overlayed from the movie in 10fps. See supplemental movie 3-4. (F) Beads migration speeds are quantified in P12 WT or *Kif6^p.G555fs^* LV wall. The violin plots show distribution with median and quartile in WT (Median=3.11, n=517frames of 53 beads on 3 LV walls) or *Kif6^p.G555fs^* (Median=1.67, n=513frames of 58 beads on 3 LV walls). P-value was determined with the Mann–Whitney test. (G) Movement tracks of cilia for 0.3 sec in the cultured ependymal cells from WT or *Kif6^p.G555fs^*, at differentiation day 14 (D14). See supplemental movie 5-7. (H) Representative kymographs of cilia in D14 cultured WT (top panel) or *Kif6^p.G555fs^* ependymal cells. *Kif6^p.G555fs^* cilia have slow motility (middle panel) or no motility (bottom panel). (I) Graph represents cilia beating frequency (Hz) in D14 cultured WT or *Kif6^p.G555fs^* ependymal cells. The truncated violin plots show distribution with median and quartile in D14 WT (Median=19.9, n=118 cilia from 3 experiments), D14 *Kif6^p.G555fs^* (Median=9.79, n=118 cilia from 3 experiments), D22 WT (Median=20.82, n=98 cilia from 2 experiments), or D14 *Kif6^p.G555fs^* (Median=23.08, n=98 cilia from 2 experiments). P-values were determined with the Mann–Whitney test. (J) Movement tracks of cilia for 0.23sec in D14 cultured ependymal cell transfected with mCherry or mCherry-Kif6 motor less (ML) as a dominant negative form of Kif6. See supplemental movie 8-9. (K) Graph indicates cilia beating frequency (Hz) in mCherry or mCherry-Kif6 ML expressing ependymal cells. The truncated violin plots show distribution with median and quartile in mCherry (Median=13.62, n=32 cilia from 3 experiments) or mCherry-Kif6 ML (Median=0.000, n=32 cilia from 3 experiments). P-value was determined with the Mann–Whitney test.

As ependymal cilia beating contributes to directional flow and CSF circulation in mouse brain, we first observed the cilia-driven flow on the surface of LV wall (Fig. 3D). Microbeads were placed on the surface of LV wall at the middle part and flowed in a linear path along the anterior-dorsal region in wild-type (WT) LV wall (Fig. 3E). On the other hand, bead displacement on *Kif6^pG555fs^* ventricles was more torturous (Fig. 3E, Movie 3-4), with speeds significantly slower than WT (Fig. 3F).

Because Kif9B (homolog of Kif6) knockdown causes motility defects in *Trypanosome* flagellum (Demonchy et al., 2009), we next asked if the *Kif6^pG555fs^* mutation also impairs ependymal cilia beating. To avoid potentially confounding effects of hydrocephalus and other environmental factors within brain ventricles, we evaluated cilia beating in primary cultured ependymal cells. Live imaging of ependymal cells showed the coordinated cilia beatings in WT and the decreased cilia motility in *Kif6^pG555fs^* (Fig. 3G, Movie 5-7). Consistent with the variable penetrance of hydrocephalus, we found at D14 that *Kif6^pG555fs^* ependymal cells had normal, faint, or no motile cilia (Fig. 3H, Movie 6) and that the beating frequency of ependymal cilia, when present, were significantly decreased in *Kif6^pG555fs^* compared with WT cilia (Fig. 3I). Interestingly, cilia beat frequency in *Kif6^pG555fs^* was not significantly different with D21 WT (Fig. 3I), suggesting that cilia beating in *Kif6^pG555fs^* was restored as ependymal cells mature.

As kinesins can form homo- and hetero-dimers, kinesins without the motor domain (motor less) can act as dominant negative proteins (Gelfand et al., 2001). To test this model, we transfected a N-terminal tagged motor-less Kif6 construct (mCherry-Kif6 ML) into ependymal cells. We observed that mCherry-Kif6 ML significantly suppressed cilia beatings in D14 ependymal cells compared with mCherry control (Fig. 3J-K, Movie 8-9). These results suggest that Kif6 regulates cilia motility in ependymal cells.

### Ependymal cilia in *Kif6^p.G555fs^* display relatively normal axoneme ultrastructure

Cilia motility requires a radially arranged nine doublet microtubules and axonemal dyneins in repeating patterns (Ma et al., 2019). To observe if ultrastructural defects may explain the decrease cilia beat frequency, we visualized cilia structure in WT and *Kif6^pG555fs^* mutant mice by Transmission Electron Microscopy (TEM). TEM images showed a typical 9+2 arrangement with outer dynein arms in both WT and *Kif6^pG555fs^* ependymal cilia at P14 (Fig. 4A). Using immunofluorescence we show that the outer dynein arm component Dnai1 was present in *Kif6^pG555fs^* ependymal cell cilia as in WT (Fig. 4B-C). Decreased cilia length is associated with decreased cilia motility (Bottier et al., 2019), however we found that the length of *Kif6^pG555fs^* ependymal cilia was not significant with WT (Fig. 4D-E). We found no structural defects in *Kif6^pG555fs^* ependymal cilia, so we turned our attention to the MCC apical cytoplasm.

**Figure 4.**
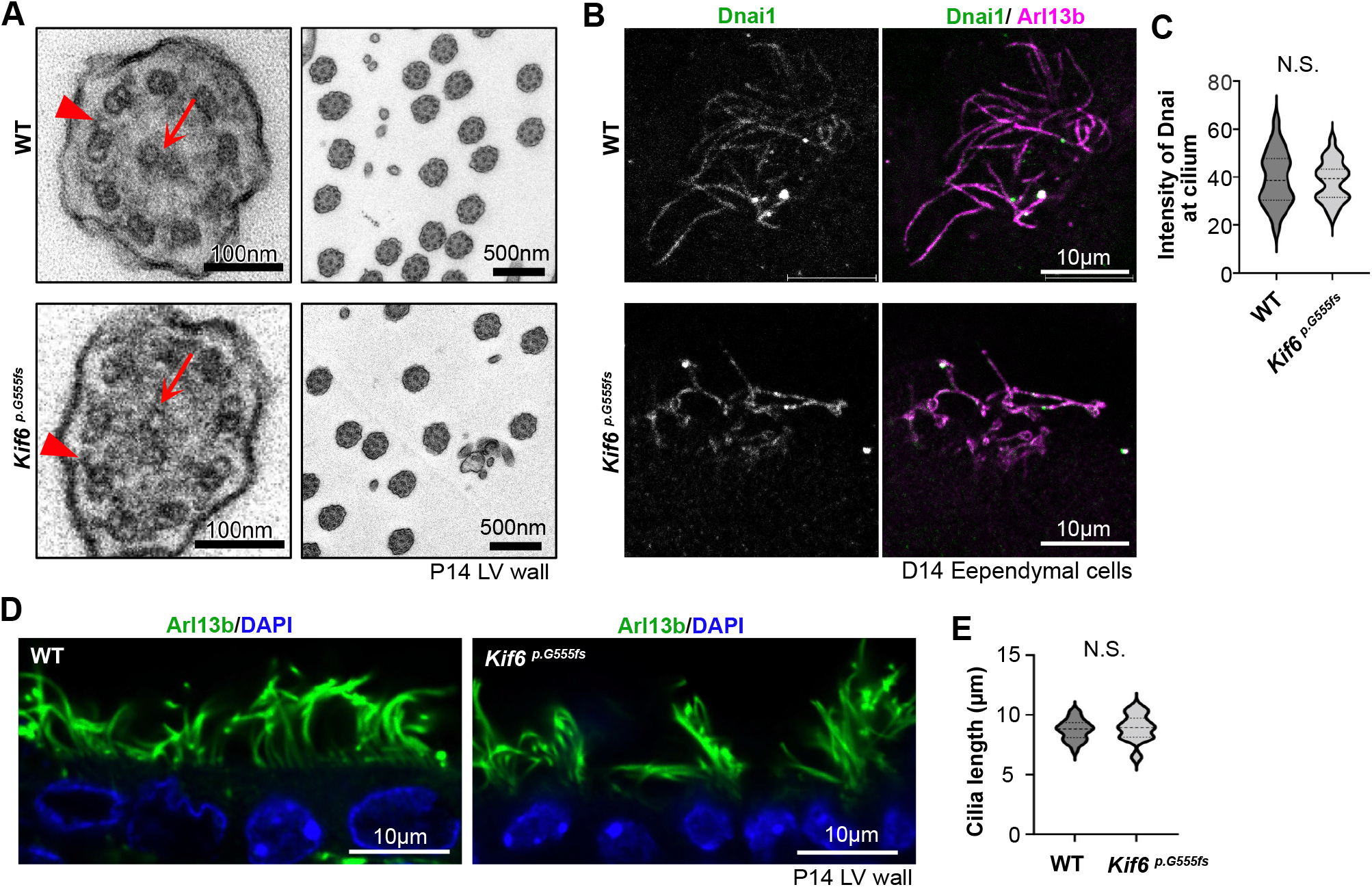
Ependymal cilia in *Kif6^p.G555fs^* display relatively normal axoneme structure. (A) Representative TEM images of ependymal cilia at P14 WT or *Kif6^p.G555fs^* LV wall. Arrows indicate central pair of microtubules. Arrowheads indicate outer dynein arm. (B) WT or *Kif6^p.G555fs^* ependymal cells at D14 were immunostained with anti-Dnai1 (green, outer dynein arm marker) and anti-Arl13b (magenta). (C) Quantification of Dnai1 intensity at cilia. Mean intensity of Dnai1 were normalized by Arl13b intensity at cilia. The violin plots show distribution with median and quartile in D14 WT (Median=38.53, n=48 cilia from three experiments) or D14 *Kif6^p.G555fs^* (Median=39.25, n=48 cilium from three experiments). P-value (0.796) was determined with the Mann Whitney test. (D) Cross section of ependymal cells in the middle lateral side of LV wall. Vibratome section from P14 WT or *Kif6^p.G555fs^* brain are immunostained with anti-Arl13b (green) and DAPI (blue). (E) Graph represents cilia length in WT or *Kif6^p.G555fs^* LV wall. The violin plots show distribution with median and quartile in P14 WT (Median=8.821, n=49 cilia from three mice) or P14 *Kif6^p.G555fs^* (Median=8.928, n=49 cilia from three mice). P-value (0.4341) was determined with the Welch’s test.

### *Kif6* is required for planar polarization of ependymal cells

Because Kif6 was localized to cytoplasmic microtubules around basal bodies (BB), we next analyzed apical microtubules and BBs in ependymal cells in *Kif6^pG555fs^* mutant mice. Apical microtubules form an organized network and align BBs in multiciliated cells (Werner et al., 2011; Kunimoto et al., 2012). Because Kif19A is known to depolymerize microtubules (Niwa et al., 2012), we first examined general microtubule organization in *Kif6^pG555fs^* ependymal cells. Overexpression of mCherry-Kif6 full length or the dominant negative Kif6 ML had no effect on acetylated-tubulin, which correlates with stable microtubules, in ependymal cell bodies (Fig. S2A-B). Moreover, *Kif6^pG555fs^* did not diminish the intensity of ß-tubulin labeled microtubule accumulation around BBs (Fig.5 A-B).

**Figure 5.**
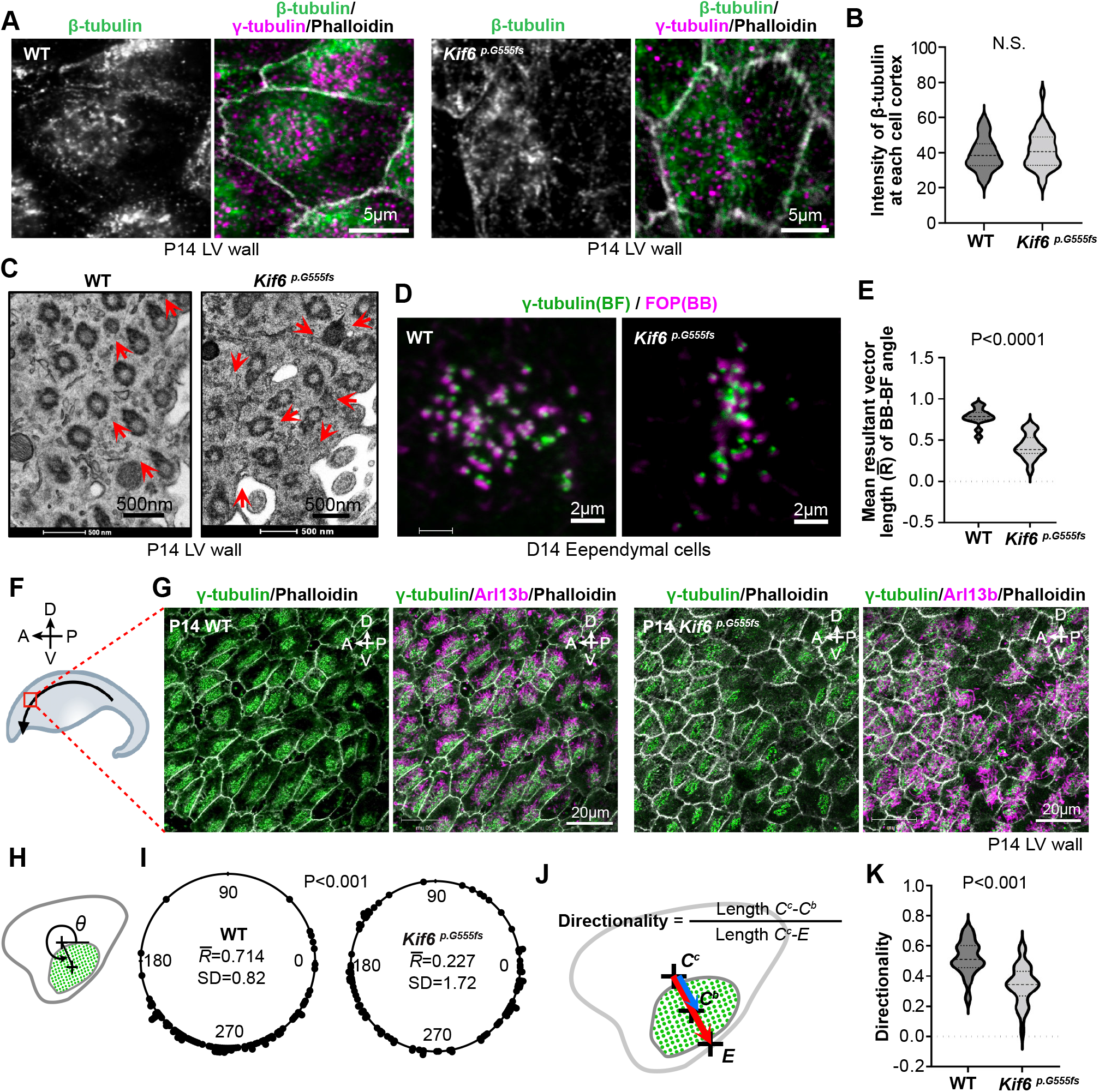
*Kif6^p.G555fs^* is required for planar polarization of ependymal cells. (A) Representative images of apical microtubules at ciliary base in P14 WT or *Kif6^p.G555fs^* ependymal cells. WT or *Kif6^p.G555fs^* LV wall are whole-mount immunostained with anti-β-tubulin (green), γ-tubulin (magenta), and Phalloidin (gray). (B) Graph represents intensity of β-tubulin at apical cell cortex as shown in (A). The violin plots show distribution with median and quartile in P14 WT (Median=38.52, n=37 cells from three mice) or P14 *Kif6^p.G555fs^* (Median=40.62, n=37 cells from three mice). P-value (0.36) was determined with the Welch’s test. (C) TEM images of ciliary base in ependymal cell from P14 WT or *Kif6^p.G555fs^* LV wall. Arrows indicate vectors from BB to BF. (D) Representative immunosataining images of γ-tubulin (green, as a BF marker) and FOP (magenta, as a BB marker) in D14 WT or *Kif6^p.G555fs^* ependymal cells. (E) Graph represents mean resultant vector length (R bar) of BB-BF angle that are measured in the immunostaining images as shown in (D). The violin plots show distribution with median and quartile in D14 WT (Median=0.79, n=15 cells from two experiments) or D14 *Kif6^p.G555fs^* (Median=0.39, n=15 cells from two experiments). P-value was determined with the Welch’s test. (F) Representation of the measurement area to observe translational polarity in LV wall tissue at P14. Whole mount-immmunostained LV walls were observed at anterior area (red box) of LV wall. Arrow indicates direction of CSF flow. (G) Representative images of whole-mount immunostaining with γ-tubulin (green) and Arl13b (magenta), and Phalloidin (gray) in the anterior area of P14 WT or *Kif6^p.G555fs^* LV walls. (H) Illustration represents the measurement of cilia bundle positioning in apical cell cortex of ependymal cell. Vector angles (θ) were calculated from center of cell to center of cilia bundle bases, which is aggregated region of γ-tubulin dots. (I) Vector angles (θ) were plotted on circular diagram in P14 WT or *Kif6^p.G555fs^* LV walls. The mean resultant vector length (R bar) and SD of θ were calculated in WT (R bar=0.714, n=100 cells from three mice) or *Kif6^p.G555fs^* (R bar=0.227, n=100 cells from three mice). P-value was determined with Watson’s two-sample test. (J) Illustration represents the measurement of “Directionality” of cilia bundle bases to cell border. The length between center of cell and center of cilia bundle bases *(C^c^-C^b^)* were divided by the length between center of cell and cell edge *(C^c^-E)*, as “Directionality”. (K) Graph represents “Directionality” of cilia bundle bases in P14 WT or *Kif6^p.G555fs^* LV walls, measured as shown in (J). The violin plots show distribution with median and quartile in P14 WT (Median=0.51, n=56 cells from three mice) or P14 *Kif6^p.G555fs^* (Median=0.34, n=56 cells from three mice). P-value was determined with the Welch’s test.

Because the surface flow was disturbed on *Kif6^pG555fs^* LV wall (Fig. 3E), we next analyzed cilia polarity/orientation at the single cell level (rotational polarity) and at the tissue level (translational polarity)(Mirzadeh et al., 2010). Rotational polarity was quantified by the orientation of the basal foot (BF) projection from BB. Using TEM of P14 LV walls, we showed that the orientation of the BF-BB was perturbed *Kif6^pG555fs^* compared to parallel orientation typically observed in WT (Fig.5C). We also observed this rotational polarity defect in cultured ependymal cells derived from *Kif6^pG555fs^* mutants by immunostaining with FGFR1 Oncogene Partner (FOP, a BB marker) and γ-tubulin (a BF marker) (Fig. 5D). The orientation of BB-BF alignments in the plane of LV tissue, was calculated as mean resultant vector lengths (R bar) and was show to be significantly reduced in *Kif6^pG555fs^* ependymal cells (Fig.5E).

Ependymal cilia also display translational polarity, visible as the clustering of BB on the LV wall along the direction of CSF flow (Ohata and Alvarez-Buylla, 2016). We observed BB clustering within each ependymal cell at the anterior-dorsal area of LV walls (Fig. 5F), finding them positioned ventrally. This translational polarity was disturbed in *Kif6^pG555fs^* (Fig. 5G). BB clustering was quantified as vector angles from the center of the cell to the center of the multicilia bundle base (Fig. 5H), which was more randomly distributed in *Kif6^pG555fs^* LV walls compared with the polarized distribution observed in WT tissues (Fig. 5I). BB clusters in *Kif6^pG555fs^* mutants were close to the center of cell (Fig. 5J) and the directionality was significantly decreased (Fig. 5K).

Finally, tyrosinated-tubulin, which marks newly polymerized microtubule plus ends, is asymmetrically localized to apical junctions in ependymal cells, and this microtubule polarization promotes cilia polarity (Takagishi et al., 2020). However, we observed no difference in tyrosinated-tubulin polarization between *Kif6^pG555fs^* and WT (Fig. S2C-D).

### Loss of Kif6 function affects apical actin assembly and BB stability

Apical actin forms networks in multiciliated cell, aiding the connection BBs and contributing to their alignment, orientation, and stabilization in multiple ciliated cells (Werner et al., 2011; Mahuzier et al., 2018). Using fluorescently labelled phalloidin, which labels actin filaments, we observed apical actin accumulation co-localized with BBs in both WT and *Kif6^pG555fs^* LV tissues (Fig. 5A). However, we observed a significant decrease in actin accumulation localized with BB clusters in *Kif6^pG555fs^* mutants (Fig. 6A, B). Moreover, the area of BB clusters was significantly increased in *Kif6^pG555fs^* ependymal cell (Fig. 6C), and were dispersed across the apical cell cortex with decreased apical actin, compared to the tight accumulation of BBs in WT ependymal cells (Fig.6A). Importantly, the loss of apical actin networks has been shown to induce BB dis-attachment in ependymal cells (Mahuzier et al., 2018). Accordingly, we found that the number of BBs was significantly decreased in *Kif6^pG555fs^* ependymal cells (Fig. 6D). Because the BBs were not found in deeper cytoplasm (Fig. 6E), these results suggest that the defect of apical actin networks in *Kif6^pG555fs^* may cause the de-stabilization of BBs and shedding of ependymal cilia.

**Figure 6.**
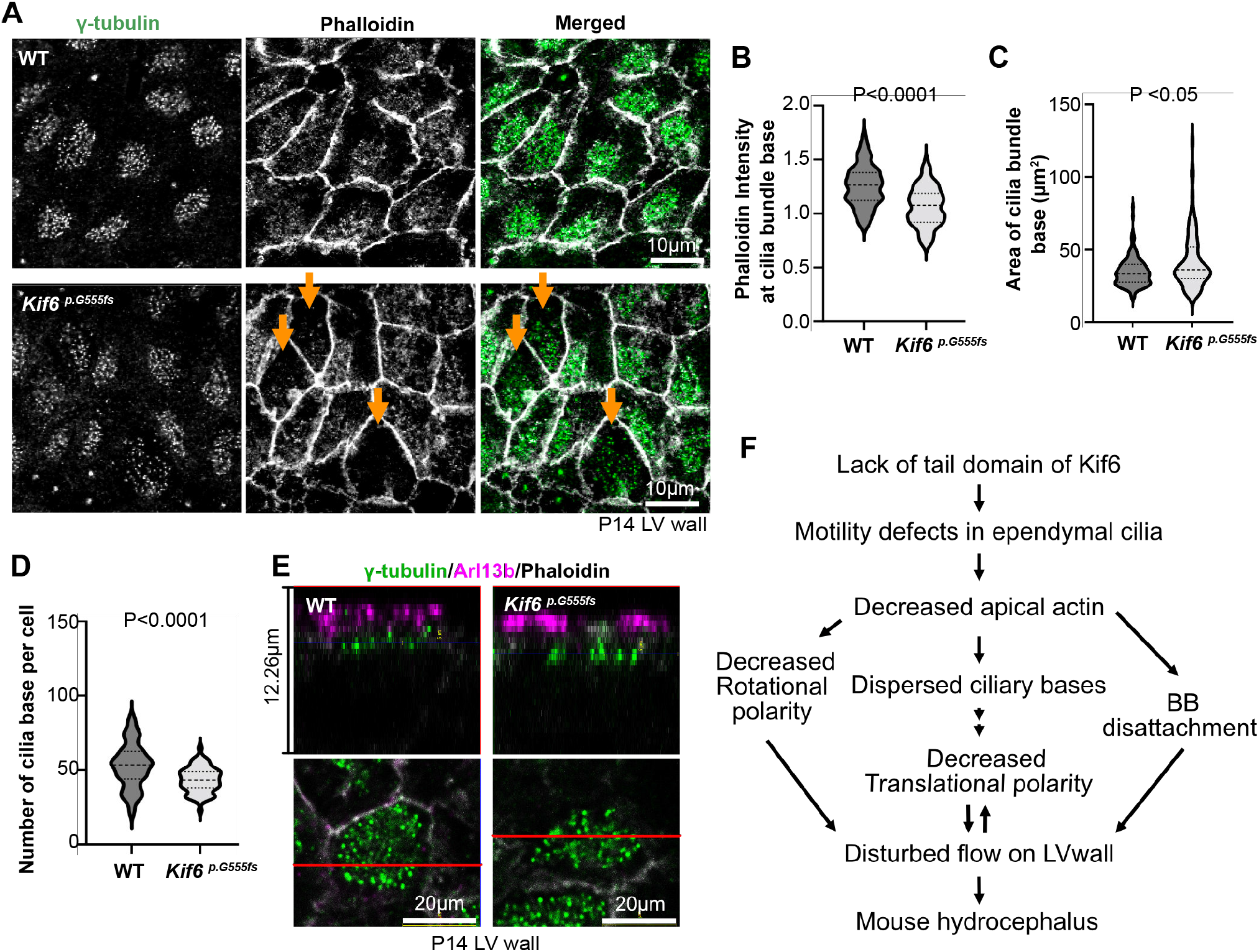
*Kif6^p.G555fs^* decreases apical actin and BB stability. (A) Representative images of whole mount immunostaining with γ-tubulin (green) and Phalloidin (gray) in the anterior area of P14 WT or *Kif6^p.G555fs^* LV walls. Arrows indicates the cells with decreased phalloidin and dispersed ciliary bases. (B) Graph represents Phalloidin intensity at cilia bundle base. The intensity of phalloidin at γ-tubulin dots area were divided by the intensity of Phalloidin at all area of apical cell cortex. The violin plots show distribution with median and quartile in P14 WT (Median=1.26, n=100 cells from three mice) or P14 *Kif6^p.G555fs^* (Median=1.08, n=99 cells from three mice). P-value was determined with the Welch’s test. (C) Graph represents the spreading area of γ-tubulin dots as the area of cilia bundle bases. The violin plots show distribution with median and quartile in P14 WT (Median=33.21, n=100 cells from three mice) or P14 *Kif6^p.G555fs^* (Median=36.08, n=101 cells from three mice). P-value (0.0162) was determined with the Mann Whitney test. (D) Graph represents the number of γ-tubulin dots in apical cell cortex as the number of cilia base per cell. The violin plots show distribution with median and quartile in P14 WT (Median=53.50, n=60 cells from three mice) or P14 *Kif6^p.G555fs^* (Median=43.5, n=60 cells from three mice). P-value was determined with the Mann Whitney test. (E) Representative confocal microscopy images of whole mount immunostaining with γ-tubulin (green), Arl13b (magenta), and Phalloidin (gray) in P14 WT or *Kif6^p.G555fs^* LV walls. Upper panels show cross sections of ependymal cell shown in lower panels at red line. (F) Effects of cilia disfunction in *Kif6^p.G555fs^* ependymal cells on mouse LV wall. Lack of tail domain of Kif6 (*Kif6^p.G555fs^*) causes motility defects in ependymal cells during brain development. Motility defects in ependymal cilia reduce apical actin network formation. Decreased apical actin results in de-stabilization of ciliary base, leading to decreased rotational polarity, dispersed ciliary bases, and BB dis-attachment, which in turn contributes to the disturbed flow on LV wall and mouse hydrocephalus.

## Discussion

Here, we describe a complex of cilia-related defects in the ependymal cells of Kif6 mutant mice. We suggest that this complex phenotype emerges from pleiotropic defects in ependymal cells. Because Kif6 actively moves along microtubules and is present in axonemes (Fig. 1), defects in cilia beating may result from the direct action of Kif6 in cilia motility. Conversely, because we demonstrate that Kif6 is required for apical actin assembly, its presence in the apical cytoplasm may explain defects in cilia polarity. Finally, we suggest that the defects in apical actin assembly may also result in the loss of cilia from ependymal cells. Together, this constellation of ciliary defects would result in defective CSF flow and moderate hydrocephalus in Kif6 mutant mice (Fig. 6F).

The role of kinesin-9 family proteins in ciliary formation and function is largely unknown. As a motile kinesin, *i.e.* a kinesin protein capable of processive movement along microtubules, Kif6 could transport components needed for axoneme beating or could itself directly influence the beating machinery. Although we were unable to observe Kif6 movement within the cilia of cultured ependymal cells, such difficulties were noted in previous work as they were unable to observe movement of the Trypanosome homologue KIF9B in flagella (Demonchy et al., 2009). It seems likely that the imaging difficulties stem from the low fluorescence signal of mKif6(1-493)-mNG in the high fluorescence background of the cellular environment. Alternatively, as suggested for Kif9 (Konjikusic et al., 2023), Kif6 may undergo microtubule-based motility along cytoplasmic microtubules and become anchored to axonemal microtubules in ependymal cell cilia. The incorporation of additional technological and/or imaging advances will be needed to distinguish these possibilities.

The motility defects in the cilia of Kif6 mutant mice may relate to the motility defects observed after Kif6 (Kif9B) knockdown in *Trypanosome*, which disrupts beating and formation of the paraflagellar rod (PFR) within the flagellar membrane (Demonchy et al., 2009). However, mammalian ependymal cilia don’t have an analogous PFR structure, so the precise locale of Kif6 action in the axoneme remains to be determined. The related kinesin family member Kif9 acts in the central pair, and loss of Kif9 disrupts cilia motility (Konjikusic et al., 2023), yet like Kif6 (Fig. 3), loss of Kif9 does not disrupt the overall axoneme architecture (Miyata et al., 2020). In fact, some redundancy between Kif6 and Kif9 might explain the finding that *Kif6^p.G555fs^* ependymal cilia display decreased motility at D14, but recovered by D21 (Fig. 1J). Another possibility is that Kif6 mutant ependymal cilia with decreased motility are physically removed (shedding) because of BB de-stabilization, as suggested by Mahuzier et al., 2018 (Fig. 6D-F), and that stable cilia that remain until D21 display normal beating.

Kif6 mutation affected not only cilia motility but also cilia polarity (Fig. 5), and these polarity defects likely contribute to the disturbed flow over LV wall (Fig. 3E). Cilia orientation is regulated by apical microtubules (Vladar et al., 2012; Taklagishi et al., 2020) and affected by apical actin networks between BBs (Werner et al., 2011), so Kif6 localization to this region is of interest as it marks a clear departure from that observed for Kif9 (Konjikusic et al., 2023). However, Kif6 mutation did not affect the overall appearance of apical microtubules (Fig. 5A-B, Fig.S2), suggesting disturbed cilia polarity might be secondary to the defects in cilia motility.

Indeed, cilia beating motility induces apical actin network organization that is crucial for BB stabilization (Mahuzier et al., 2018). Suppressed cilia motility would be expected to result in decreased apical actin networks around BBs, and de-stabilized BBs could be dis-attached (Mahuzier et al., 2018). Because we observed all of these phenotypes, it is possible that despite the presence of Kif6 in the apical cytoplasm, that Kif6-related actin defects are secondary to cilia beating defects.

## Materials and Methods

### Mouse experiments

*Kif6^pG555fs^* mice were generated using CRISPR-Cas9 system (Konjikusic et al., 2018). *Kif6^pG555fs^* or C57Bl/6J mice were used for whole-mount staining, ependymal cell culture, and LV explants cultures. All mice were maintained with 12/12 h light/dark cycle and no more than 5 mice per cage. Both male and female mice were used for the experiments, and sex differences were not observed. All mouse experiments were approved by the Animal Studies Committee at University of Texas at Austin (AUP-2021-00118).

### Plasmids

Mouse Kif6 cDNA encodes 803 amino acid protein (NP_796026) was cloned from a mouse brain cDNA library and sequenced. The cDNAs of mKif6 FL (1-803aa) and ML (360-803aa) were subcloned into pmCherry2-C1 vector (a gift from Michael Davidson, Addgene plasmid # 54563). mCherry was removed and mKif6 (1-493aa)-mNeonGreen was added into pmCherry2 vector. pCS2-dKif6 (1-501)-EGFP was generated from the pCS2-dKif6-EGFP plasmid by Gibson assembly (NEB HiFi DNA Assembly Kit) using a GeneArt gene fragment encoding amino acids 1-501 (Thermo Fisher).

### Cell culture, transfection and lysate preparation

COS-7 (monkey kidney fibroblast) cells obtained from ATCC (RRID: CVCL_0224) were cultured in DMEM (Gibco) with 10% (vol/vol) Fetal Clone III (HyClone) and 1% GlutaMAX (Gibco) at 37°C with 5% CO_2_. COS-7 cells were transfected with Trans-IT LT1 (Mirus) according to the manufacturer’s instructions.

COS-7 cells were collected 48 h post-transfection. The cells were harvested by low-speed centrifugation at 1,500 × g for 5 min at 4°C. The pellet was rinsed once in PBS and resuspended in ice-cold lysis buffer (25 mM HEPES/KOH, 115 mM potassium acetate, 5 mM sodium acetate, 5 mM MgCl_2_, 0.5 mM EGTA, and 1% Triton X-100, pH 7.4) freshly supplemented with 1 mM ATP, 1 mM phenylmethylsulfonyl fluoride, and protease inhibitors (P8340; Sigma-Aldrich). After the lysate was clarified by centrifugation at 20,000 × g for 10 min at 4°C, aliquots of the supernatant were snap-frozen in liquid nitrogen and stored at −80°C until further use.

### Single-molecule motility assays

HiLyte647-labeled microtubules were polymerized from purified tubulin including 10% Hily647-labeled tubulin (Cytoskeleton) in BRB80 buffer (80 mM Pipes/KOH pH 6.8, 1 mM MgCl_2_, and 1 mM EGTA) supplemented with 1 mM GTP and 2.5 mM MgCl_2_ at 37°C for 30 min. 20 μM taxol in prewarmed BRB80 buffer was added and incubated at 37°C for additional 30 min to stabilize microtubules. Microtubules were stored in the dark at room temperature for further use.

A flow cell (~10 μl volume) was assembled by attaching a clean #1.5 coverslip (Fisher Scientific) to a glass slide (Fisher Scientific) with two strips of double-sided tape. Polymerized microtubules were diluted in BRB80 buffer supplemented with 10 μM taxol and then were infused into flow cells and incubated for 5 min at room temperature for nonspecific adsorption to the coverslips. Subsequently, blocking buffer [15 mg/ml BSA in P12 buffer (12 mM Pipes/KOH pH 6.8, 1 mM MgCl_2_, and 1 mM EGTA)] was infused and incubated for 5 min. Finally, KIF6 cell lysate in the motility mixture [2 mM ATP, 0.4 mg/ml casein, 6 mg/ml BSA, 10 μM taxol, and oxygen scavenging (1 mM DTT, 1 mM MgCl_2_, 10 mM glucose, 0.2 mg/ml glucose oxidase, and 0.08 mg/ml catalase) in P12 buffer] was added to the flow cells. The flow-cell was sealed with molten paraffin wax.

Images were acquired by TIRF microscopy using an inverted microscope Ti-E/B equipped with the perfect focus system (Nikon), a 100× 1.49 NA oil immersion TIRF objective (Nikon), three 20-mW diode lasers (488 nm, 561 nm, and 640 nm) and an electron-multiplying charge-coupled device detector (iXon X3DU897; Andor Technology). Image acquisition was controlled using Nikon Elements software and all assays were performed at room temperature. Images were acquired continuously every 200 s for 40 s.

Maximum-intensity projections were generated, and the kymographs were produced by drawing along tracks of motors (width= 3 pixels) using Fiji/ImageJ2. Motors frequently paused during motility events, and thus the velocity between pauses were analyzed. Velocity was defined as the distance on the x axis of the kymograph divided by the time on the y axis of the kymograph.

### Ependymal cell culture

LV walls from P0-1 mouse brains were collected and the suspended cells with trypsin-EDTA were cultured in DMEM with 10% FBS for 3 days on a laminin-coated flask. After shaking 2hr, the adherent cells were plated on poly-L-lysine and laminin-coated coverslips and differentiated into ciliated ependymal cells (ependymal cells) by serum starvation. ependymal cells were transfected with mCherry-Kif6 constructs at 48hr before experiments using Lipofectamine 2000 (Invitrogen).

### Live cell imaging

For live imaging of Kif6 (1-493)-mNG, D5 ependymal cells were transfected with Kif6 (1-493)-mNG imaged at D7 for 2 sec in 50 fps, using Ti2E microscope with high content screening system (Nikon). To observe cilia beatings, ependymal cells were plated on the glass-bottom dish. Time-lapse images of DIC were collected for 2 sec in 47, 95 or 190 fps, using Ti2E microscope with high content screening system (Nikon). Ciliary tips were tracked by manual tracking of Fiji for images in Fig. 3G and 3J. Kymographs were generated along the track of ciliary tip and used for the measurement of cilia beating frequency.

### Western blotting

LV wall tissues were collected from 3 mice at P0, 4, 7, or 14. LV wall tissues or P45 testis tissue were lysed in PHEM buffer (50 mM PIPES, 50 mM HEPES, 1 mM EDTA, 2 mM MgSO4, 1% Triton X-100) with protease inhibitors, homogenized, and centrifuged for 1 hr at 14,000 rpm. The lysate supernatant was mixed with Laemmli Sample buffer (Bio-Rad) and 2-mercaptoehanol and boiled. Samples were separated by SDS-PAGE and transferred to Nitrocellulose membranes. The membranes were blocked with 2% BSA and probed with anti-Kif6 (1:1000, Proteintech, 17290-I-AP) or anti-GAPDH (1:1000, Cell Signaling Technology, #2118) antibodies. Detection was carried out using HRP-linked anti-Rabbit IgG (1:2000, Cell signaling Technology, #7074) and visualized using Pierce ECL substrate (Thermo Fisher).

### H&E staining and immunostaining

Mouse brains were fixed in 4% paraformaldehyde at 4°C for overnight. The fixed whole brains were paraffin-embedded and cut into 5μm sections for H&E staining. To measure cilia length, The fixed brains were sectioned in the coronal plane at 100 μm on a vibratome and immunostained. Whole-mount preparations of LV wall tissue were fixed with cold methanol for 10min and 4% paraformaldehyde for overnight at 4°C. Cultured ependymal cells were fixed with cold methanol for 3min and 4% paraformaldehyde for 10 min at RT. Whole-mount, ependymal cell, or vibratome sectioned samples were blocked with 10% goat serum in PBS for 30 min and incubated with primary antibodies for overnight at 4°C. Primary antibodies, which were anti-Kif6 (1:100, Proteintech, 17290-I-AP), anti-Arl13b (1:200, Proteintech, 17711-I-AP), anti-Acetylated Tubulin (1:200, Sigma, T7451), anti-gamma Tubulin (1:100, Abcam, ab11321), anti-beta Tubulin (1:100, Abcam, ab7287), anti-Dnai1 (1:100, NeuroMab, 73-372), or anti-Tubulin [YL 1/2] (1:200, Abcam, ab6160), were detected with secondary antibodies conjugated to Alexa Fluorophores (Invitrogen). The apical surface of ependymal cells was imaged using a confocal laser scanning microscope LSM700 (Zeiss) or LSM980 Airyscan2 SR mode with joint Deconvolution (Nikon) for super-resolution images. H&E or immune-stained tissue sections were imaged by BZ-X710 (KEYENCE).

### Beads migration on LV wall

Lateral side of LV walls were dissected from P12 mouse brain and pinned on a silicon plate with DMEM. Fluorescent microsphere (1μm) containing PBS was placed on the middle part of LV wall. The beads migration was recorded in 10 fps for 10sec by AXIO Zoom.V16 (Zeiss). Time projection images were created from 100 frames by Fiji software (Schindelin et al., 2012). Beads migration speed was calculated using the manual tracking plug-in of Fiji.

### Electron microscopy

P14 LV wall tissues were fixed using 3% glutaraldehyde with 2% paraformaldehyde for overnight. The tissues were post-fixed in 1% osmium tetroxide with 1% potassium ferrocyanide for 3hr. After en-bloc stain with 1% aqueous uranyl acetate for 1hr, ethanol-dehydrated tissues were displaced into aceton and embedded in an epoxy resin. Tissues were sectioned (80 nm thickness) with UltraCut Ultramicrotome (Leica). Images of the apical cell cortex of ependymal cells were obtained using Tecnai Transmission Electron Microscope (FEI).

### Statistical analyses

The violin plots were generated using GraphPad Prism 9 (GraphPad) Welch’s t-test or Mann-Whitney U test was used to compare the means of two experimental groups using GraphPad Prism. Differences were considered statistically significant at P < 0.05. To determine the rotational BB orientation for a single ependymal cell (Fig.5D-E), a vector was drawn from the center of FOP dot to the center of the closest γ-tubulin dot in each cell, and more than 10 vectors were averaged. Vector angles were measured using Fiji and plotted on a circular diagram using the statistical software R (R Core Team, 2019). Mean resultant vector length (R bar) were calculated using the “circular” package (Lund and Agostinelli, 2011) within the R statistical computing environment. For Fig.5I, Distribution of vector angles of cilia bundle on WT and *Kif6^pG555fs^* LV walls were compared using Watson’s two-sample U2 test by R software.

## Acknowledgements

Kif6 mice was generated by Mia J. Konjikusic (University of California, San Francisco) and administrated by Elle C. Roberson (CU Anschutz, Colorado). Live imaging of cilia beatings and super-resolution imaging were performed at Microscopy and Imaging Facility at UT Austin (RRID# SCR_021756).

## Supplementary information

**Movie 1. Movement of GFP-dKif6 (1-501) along taxol-stabilized microtubules *in vitro*, shown in Fig. 1C**

Cell lysates were collected from COS-7 cells expressing GFP-dKif6 (1-501) and added to a flow cell containing Alexa647-labeled-taxol-stabilized microtubules. Time-lapse imaging was acquired at 5 frame/sec (fps) for 40 sec. The display rate of movie is 60 fps.

**Movie 2. Live imaging of mKif6 (1-493)-mNG at a developing ependymal cilium, shown in Fig. 1G**

mKif6 (1-493)-mNG was transfected in D5 ependymal cell, and mKif6 (1-493)-mNG at short developing cilia was imaged at D7. Time-lapse imaging was acquired at 50 fps for 20 sec. Notes: the upper dot is a tip of another cilium.

**Movie 3. Beads migration on WT LVW, shown in Fig. 3E**

Fluorescent microsphere was placed on D12 WT LV wall. The beads migration was recorded in 10 fps for 10sec.

**Movie 4. Beads migration on *Kif6^p0G555fs^* LVW, shown in Fig. 3E**

Fluorescent microsphere was placed on D12 *Kif6^p.G555fs^* LV wall. The beads migration was recorded in 10 fps for 10sec.

**Movie 5. Live imaging of WT ependymal cell cilia at D14**

Primary cultured ependymal cells form WT LV walls were cultured on glass-bottom dish. Time-lapse imaging of DIC was acquired at 47 fps for 2 sec.

**Movie 6. Live imaging of *Kif6^p.G555fs^* ependymal cell cilia at D14**

Primary cultured ependymal cells form *Kif6^p.G555fs^* LV walls were cultured on glass-bottom dish. Time-lapse imaging of DIC was acquired at 47 fps for 2 sec.

**Movie 7. Live imaging of WT (left) and *Kif6^p.G555fs^* (right) ependymal cell cilia at D14, shown in Fig. 3G**

Primary cultured ependymal cells form WT or *Kif6^p.G555fs^* LV walls were cultured on glass-bottom dishes. Time-lapse imaging of DIC was acquired at 95 fps for 2 sec.

**Movie 8. Live imaging of mCherry-expressing ependymal cell cilia at D14, related to Fig. 3J**

Primary cultured ependymal cells form WT LV walls were cultured on glass-bottom dishes and transfected with mCherry at D12. At D14, time-lapse imaging of DIC was acquired at 190 fps for 2 sec. mCherry signal with DIC image is shown in the first flame.

**Movie 9. Live imaging of mCherry-Kif6-ML-expressing ependymal cell cilia at D14, related to Fig. 3J**

Primary cultured ependymal cells form WT LV walls were cultured on glass-bottom dishes and transfected with mCherry-Kif6-ML at D12. At D14, time-lapse imaging of DIC was acquired at 190 fps for 2 sec. mCherry signal with DIC image is shown in the first flame.

---

**Figure S1.**
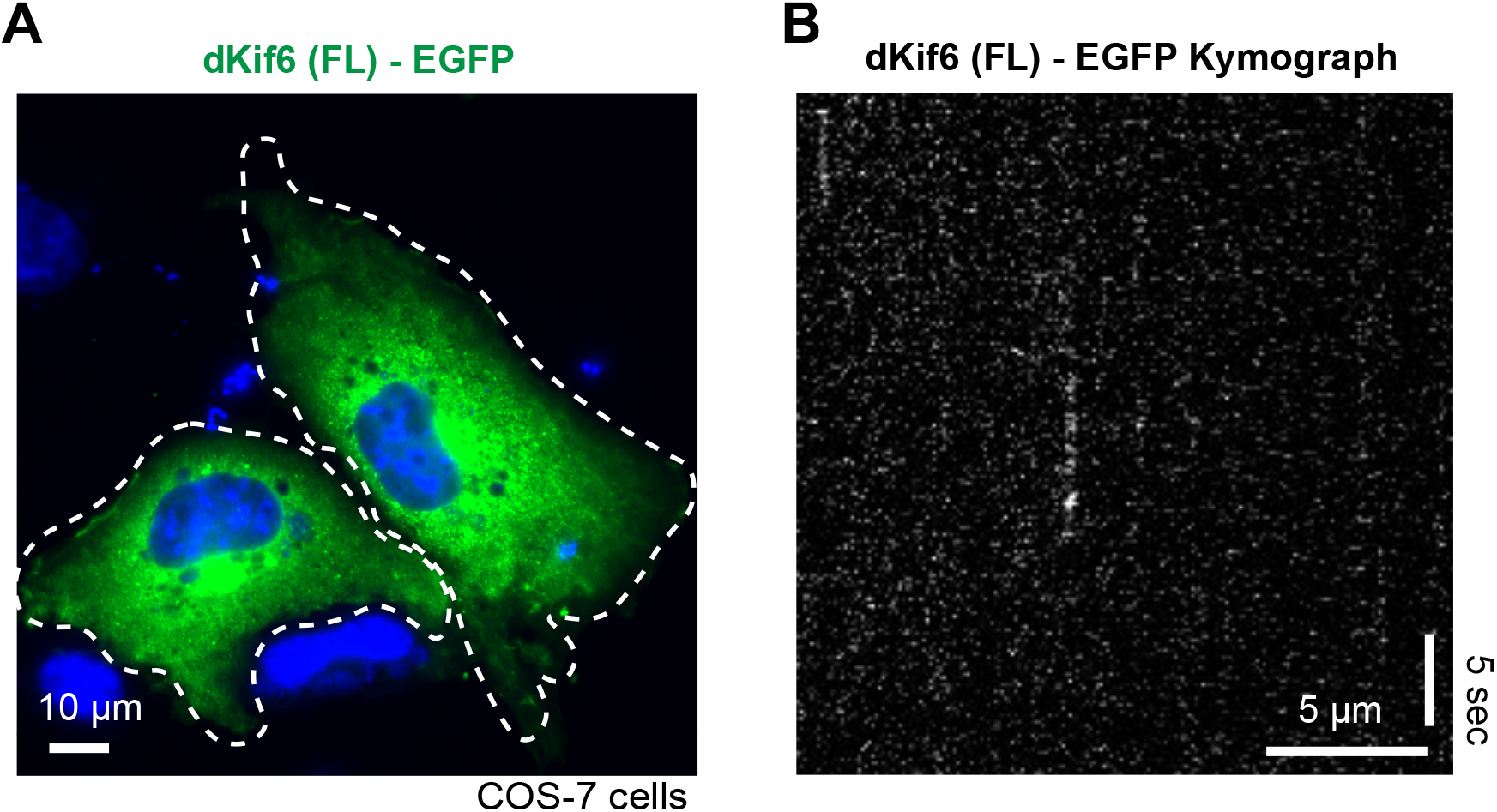
Full-length dKif6 is regulated by autoinhibition. (A) Representative image of GFP-tagged dKif6(FL) expressed in COS-7 cells. White dashed line indicates the outline of a transfected cell. Blue, DAPI stain. (B) Representative kymograph from single-molecule motility assay of dKif6(FL)-EGFP molecules on taxol-stabilized microtubules. No binding or movement of dKif6(FL)-EGFP was observed. Time is on the y-axis (scale bar, 5 sec) and distance is on the x-axis (scale bar, 5 mm).

**Figure S2.**
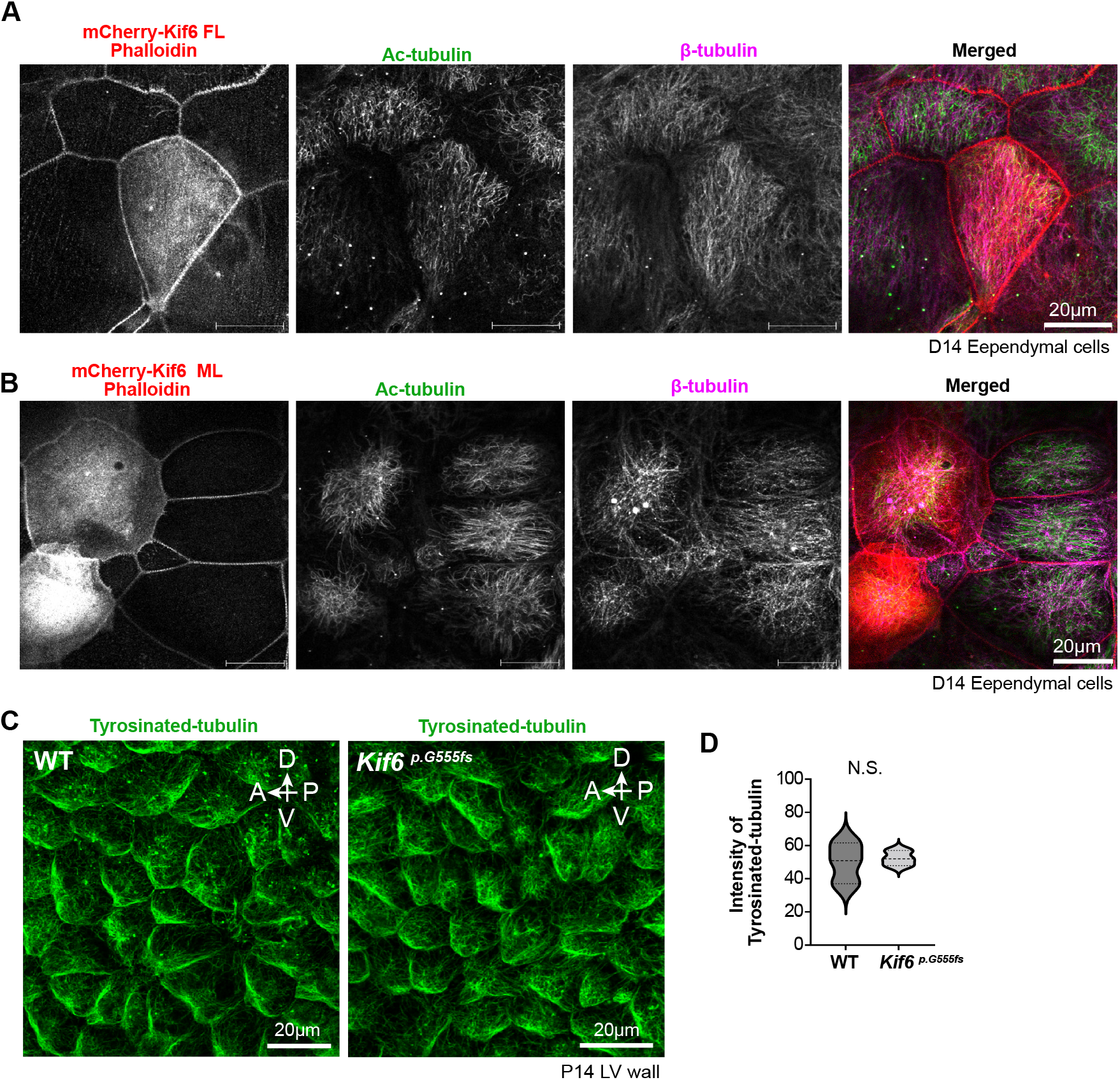
Kif6 did not affect overall appearance of microtubules, related to Fig.5. (A, B) Representative images of immunostaining with Ac-tubulin (green) and β-tubulin (magenta) in mCherry-Kif6 FL (A) or mCherry-Kif6 ML (B) expressing D14 ependymal cell. (C) Representative images of whole-mount immunostaining with Tyrosinated-tubulin (green) in P14 WT or in *Kif6^p.G555fs^* LV wall at the apical cell cortex of anterior area. (D) Graph represents the intensity of Tyrosinated-tubulin at apical cell cortex as shown in (C). The violin plots show distribution with median and quartile in P14 WT (Median=50.93, n=3 mice) or P14 Kif6p.G555fs (Median=52.01, n=3 cells mice). P-value (0.7717) was determined with Welch’s test.

